# The length of the thalamo-cortical white matter fibers brings insight into sex differences in sleep spindle frequency

**DOI:** 10.1101/2022.05.11.491489

**Authors:** Pierre-Olivier Gaudreault, Jean-Marc Lina, Maxime Descoteaux, Nadia Gosselin, Julien Doyon, Samuel Deslauriers-Gauthier, Julie Carrier

**Affiliations:** Center for Advanced Research in Sleep Medicine, CIUSSS du Nord-de-l’île-de-Montréal -Hôpital du Sacré-Cœur de Montréal, Montreal, Que., Canada; Department of Psychiatry, Icahn School of Medicine at Mount Sinai, New York City, NY, USA; Department of Electrical Engineering, École de Technologie Supérieure, Montreal, Que., Canada; Sherbrooke Connectivity Imaging Lab, Computer Science Department, Université de Sherbrooke, Sherbrooke, Que., Canada; Department of Psychology, Université de Montréal, Montreal, Que., Canada; Research Center, Institut Universitaire de Gériatrie de Montréal, Montreal, Que., Canada; McConnell Brain Imaging Center, Montreal Neurological Institute, McGill University, Montreal, Que., Canada; Inria Sophia Antipolis Méditerranée Research Center, Université Côte d’Azur, France

**Author notes:** **Corresponding author:** Julie Carrier, PhD, Center for Advanced Research in Sleep Medicine, Research Center - Hôpital du Sacré-Cœur de Montréal, 5400 Gouin West Blvd. Montreal, Quebec, H4J 1C5, Canada. Both S.D-G and J.C. contributed equally to the supervision of the study as senior researchers.

**Keywords:** Thalamo-cortical loop, sex differences, diffusion MRI, fiber bundle length

## Abstract

Sleep spindles (SS) are crucial to brain functions like memory and learning. SS characteristics result from the propagation of nerve impulses along white matter (WM) projections underlying an intricate loop between the thalamus and the cortex. SS amplitude and density have been associated with WM diffusion microarchitecture but physiological mechanisms underlying individual and sex-related variations in SS frequency are unknown. Here, we tested a model of traveling signals along the thalamo-cortico-thalamic projections to explain individual differences in spindle frequency. We predicted the presence of a relationship between the length of the thalamo-cortical WM bundles and a specific characteristic of this functional network, SS frequency.

Thirty young participants underwent a polysomnographic recording and a 3T MRI including a diffusion sequence. The length of WM fiber bundles between the thalamus and the frontal cortex was derived from probabilistic tractography computed through constrained spherical deconvolution.

Longer WM fiber bundles between the thalamus and specific regions of the frontal cortex (rostral middle frontal gyrus and anterior and middle part of the superior frontal gyrus) were associated with slower SS frequency. Moreover, the length of these WM fiber bundles statistically mediated the sex-related differences in SS frequency.

By providing a neuroanatomical marker of individual and sex-related differences in SS frequency, this study is the first to highlight the association between the anatomy of a specific brain network and a specific functional characteristic of this network, the frequency of oscillations produced during sleep.

## Introduction

Among the characteristics of brain oscillations visible on the electroencephalographic (EEG) recordings, the intrinsic frequency of rhythmic transients is a fundamental property associated with specific cognitive functions (Kahana, 2006). Sleep spindles (SS), an oscillatory hallmark of non-rapid-eye-movement (NREM) sleep, are short bursts (≈0.7s) of fusiform oscillatory brain activity (demonstrating waxing and waning phases) between 12 and 16 Hz observed predominantly in the centro-parietal regions of the scalp (De Gennaro and Ferrara, 2003). Neuronal activity during SS has been implicated in the modulation of neural responses to stimuli, decreasing the brain’s permeability to external stimuli according to their salience in order to ensure sleep maintenance and protection (Blume et al., 2018). Notably, SS density (number of spindles per min of NREM sleep), amplitude, and oscillatory frequency are also associated with neuronal plasticity processes, cognitive integrity, and long-term brain functionality including declarative and procedural sleep-dependent memory consolidation (see Fogel and Smith, 2011 for reviews). SS are also associated with individual differences in global cognitive abilities including fluid intelligence measures and the intelligence quotient (IQ) (Fang et al., 2017; Ujma et al., 2014).

SS show substantial individual variability but are relatively consistent from one night to another for one individual, making them a stringent EEG signature that is unique for each individual (Cox et al., 2017). Most importantly, characteristics of SS constitute one of the most robust sex differences in the sleeping EEG (See Carrier et al., 2017 for review). For instance, women show higher NREM sleep EEG spectral power in the SS-related sigma frequency band (12-16 Hz), higher SS density, greater SS amplitude, and faster SS frequency as compared to men (Carrier et al., 2001; Martin et al., 2013; Ujma et al., 2014). The origins of those sex differences are currently unknown although some possible mechanisms encompassing sex steroid hormones have been postulated since SS are known to vary during pregnancy and across the menstrual cycle (Carrier et al., 2017). SS are generated via a complex activation of the thalamo-cortical loop (see Clawson et al., 2016 for a review). Activation of the GABAergic neurons of the thalamic reticular nucleus causes highly synchronized and rhythmic bursts of inhibitory firing in thalamo-cortical neurons mainly in dorsal thalamic nuclei. Post-inhibitory rebound activity in thalamo-cortical neurons generates glutamatergic excitatory potentials in both the cortex and the thalamic reticular nucleus, preparing them for the next firing burst (Steriade, 2000). Although thalamic nuclei are central to SS generation, the entire thalamo-cortico-thalamic loop is necessary for the synchronization and termination of the SS observed on the scalp EEG. Indeed, as successive volleys of excitatory thalamo-cortical input increase cortical activity, they also increase cortico-thalamic feedback to the thalamic reticular nucleus and dorsal thalamic nuclei. This excitatory feedback synchronizes successive cycles of activity between thalamic reticular nucleus and thalamo-cortical neurons (Lüthi, 2014), but also increases the firing rate in dorsal thalamic nuclei which desynchronizes the firing of cortico-thalamic projections. Ultimately, this disruption of intrathalamic oscillations causes SS waning (Bonjean et al., 2011; Timofeev et al., 2001).

This complex dialog between the thalamus and the cortex partly relies on axonal projections. Diffusion magnetic resonance imaging (dMRI), a technique assessing the diffusion of water molecules in the brain, is considered a sensitive measure of the underlying white matter (WM) microstructure through metrics representing the coherence of diffusivity along WM fiber orientations. Two studies used dMRI in a whole-brain voxelwise approach to assess the link between individual SS variables and WM diffusion metrics. These metrics were found to significantly predict SS density, amplitude, and power, as well as sigma spectral power (Gaudreault et al., 2018; Piantoni et al., 2013). It was more specifically hypothesized that frontal WM integrity underlying the thalamo-cortical loop is a good predictor of SS density and amplitude in healthy young individuals by enhancing neuronal synchrony of the underlying neurons and ultimately impacting the EEG (Gaudreault et al., 2018). However, no association was reported between SS mean frequency and WM microstructure.

SS characteristics measured on the scalp result from the closed-loop propagation of neural impulses between the thalamus and the cerebral cortex (Mak-McCully et al., 2017), representing the back-and-forth communication along WM projections. This model of propagation originated from landmark electrophysiological studies that first demonstrated the intrathalamic generation of spindling oscillations, but also the recruitment of cortical circuits through rhythmic synaptic currents in phase with thalamic activity (See Fernandez and Lüthi, 2020 for an extensive review). The cortical state of a SS can therefore be considered as being generated in the cortical layers through a traveling wave from the thalamus and back with a specific velocity (*v*), an oscillating frequency (*f*), and a wavelength (λ) that corresponds to the spatial periodicity of the instantaneous nerve impulse profile along the thalamo-cortical projections. The largest and fundamental wavelength corresponds to the total length of the WM fibers involved in the thalamo-cortical loop. As illustrated in Figure 1, for a given propagation speed *v* of neural impulses, the frequency *f* of a SS measured locally on the cortex must be related to the wavelength λ through the relation *v*/*f* = *λ*. Assuming those traveling waves synchronize the cortical generators to where the thalamo-cortical fibers project, the SS frequency *f* should be related to the inverse of the wavelength λ that is proportional to the distance separating the thalamus from the cerebral cortex. Indeed, this specific relationship between brain rhythms and neurophysiological factors has been confirmed through computational modeling (Robinson et al., 2001). In terms of physiology, the action potentials generated by the glutamatergic post-inhibitory rebound activity of the thalamo-cortical neurons should reach cortico-thalamic neurons more quickly in shorter WM projections. By rapidly reaching cortico-thalamic neurons, the excitatory post-synaptic potentials would increase the probability of depolarizing cortico-thalamic neurons that ultimately send their inputs back to the thalamus in order to keep intrathalamic synchrony and prepare the system for the next volley of burst firing (Clawson et al., 2016; Lüthi, 2014). The time needed for the reciprocal neuronal firing between thalamo-cortical and cortico-thalamic neurons along shorter WM projections should increase the number of complete cycles in the thalamo-cortical loop, and in turn increase the SS frequency.

**Figure 1.**
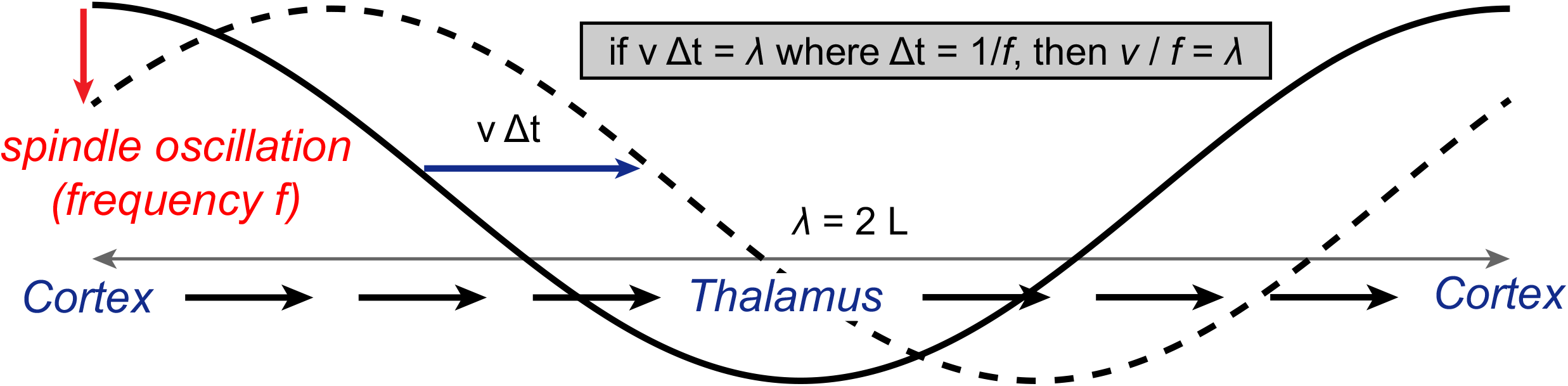
Schematic representation of the traveling wave in the generating of sleep spindles. Propagation of an oscillatory signal as a traveling wave with a speed *v* (m/s), during Δt seconds, along the thalamo-cortical loop (length λ = 2L). During this propagation, the cortical signal, i.e., the *spindling* activity oscillates with frequency *f*. After one period, this frequency measured locally on the cortex must be related to the length λ through the relation *v*/*f* = *λ*.

In addition to fiber bundle length, multiple factors have been shown to impact conduction velocity, including myelin thickness, internodal spacing, and most importantly axon diameter (Caminiti et al., 2013; Drakesmith et al., 2019). Variations of such factors could then affect the frequency of spindling oscillations. The adequate estimation of most of these parameters *in vivo*, however, needs further methodological advancements and therefore led us to use a quantifiable measure derived from diffusion MRI, the specific length of the axonal projections between the thalamus and regions of the frontal cortex. Our study aimed at testing this propagation model in healthy adults through the relationship between the anatomy of the thalamo-cortical loop and a specific functional characteristic of this network, the SS frequency. By specifically targeting the frontal and central regions of the cortex, we hypothesized that longer streamlines between the thalamic nuclei and these anterior regions of interest (ROIs) will be associated with slower SS frequency. We proposed the use of a streamline-based and bundle-specific approach in order to quantify streamlines between the thalamus and the frontal cortex. Probabilistic tractography was performed by taking anatomical constraints into account, leading to more robust WM bundle identification. We further tested whether sex differences in SS frequency could statistically be explained by the length of WM bundles. We also hypothesized that faster SS mean frequency in women would statistically be explained by sex differences in WM fiber length when correcting for the intracranial volume (ICV). The present work makes, for the first time, a bridge between the anatomy of a specific network of the human brain and the frequency of a sleep-specific functional oscillation, the sleep spindles.

## Materials and Methods

### Participants

Thirty young adults between 20 and 30 years of age (mean: 22.9 ± 2.8 yo; 14 women) completed the research protocol. Extensive exclusion criteria were used to minimize factors which may impact sleep measures. The related information was collected through an in-house questionnaire, and a semi-structured interview. Exclusion criteria included smoking, a body mass index (BMI) >27, the use of drugs or medication known to affect the sleep-wake cycle or the nervous system, any complaints about the sleep-wake cycle, a habitual sleep duration <7h or >9h, night shifts, or transmeridian travels within three months prior to the study, and any history of neurological or psychiatric illness. Participants with subclinical scores (>13) on the Beck Depression Inventory and (>7) on the Beck Anxiety Inventory were excluded. Participants also underwent a complete neuropsychological evaluation to rule out major cognitive deficits. Each participant underwent two nights of polysomnographic (PSG) recording at the laboratory: one adaptation/screening night and one experimental night. The adaptation/screening night included EEG, leg electromyogram (EMG), thoracoabdominal plethysmograph, and oral/nasal cannula. Participants were excluded if they showed an apnea/hypopnea index >10 per hour of sleep or an index of periodic leg movements associated with microarousals >10. The research protocol was approved by the Hôpital du Sacré-Coeur de Montréal Institutional Review Board and the Unité de neuroimagerie fonctionnelle (UNF) research ethics mixed committee. Written consent was obtained from all subjects prior to the study and monetary compensation was given to each for their participation.

### Polysomnographic recordings and spindle detection

All subjects underwent a night of PSG recording after the screening night. Electroencephalography, chin electromyogram, electrooculogram, and electrocardiogram signals were recorded on a Grass Model 15A54 amplifier system (Natus Neurology, Warwick, Rhode Island, USA). EEG included 20 EEG derivations (Fp1, Fp2, Fz, F3, F4, F7, F8, Cz, C3, C4, Pz, P3, P4, Oz, O1, O2, T3, T4, T5, T6) referred to linked earlobes (10-20 international system; bandpass 0.3–100 Hz; sampling rate of 256 Hz) and the digitalization of signals was carried out using a commercial software (Harmonie, Stellate Systems, Montreal, Quebec, Canada). Sleep stages (N1, N2, N3, and REM) were visually scored by a trained sleep technician according to standard AASM criteria (Iber et al., 2007) using 30-second epochs. Artefacts were automatically detected as described elsewhere (Gaudreault et al., 2018) and visually confirmed by a trained technician. SS were automatically detected using a previously published detection algorithm (Gaudreault et al., 2018; Martin et al., 2013) on artefact-free NREM epochs in parasagittal derivations (Fp1, Fp2, F3, F4, Fz, C3, C4, Cz, P3, P4 and Pz). First, EEG data were filtered with a linear phase finite impulse response filter (–3 dB at 11 and 15 Hz). Zero-phase distortion was obtained by filtering the data forward and reverse. Then, using a 0.25-second time window, the root mean square of the filtered signal was calculated and thresholded at the 95th percentile. A criterion of at least two consecutive root mean square time points exceeding the threshold duration (0.5 s) was used to detect a SS. Analyzes were carried out on SS detected in NREM sleep on F3, F4, C3, and C4 derivations since the associated scalp localization corroborates the targeted WM tracts. The averaged SS frequency (in Hz) and SS amplitude (expressed in μV) were then calculated for the entire sleep period.

### MRI acquisitions

All MRI data acquisition were performed at the Unité de Neuroimagerie Fonctionnelle (UNF) at the Research Center of the Institut Universitaire de Gériatrie de Montréal using a 3T Siemens Trio MRI scanner (Siemens Medical Systems, Erlengan, Germany). The MRI diffusion sequence included an echo planar imaging (EPI) sequence with the following parameters: repetition time TR = 12700 ms, echo time (TE) = 100 ms, bandwidth = 1302 Hz/Px; 128 × 128 acquisition matrix, 75 slices, antero-posterior acquisition; one reference image at b = 0 s/mm^2^ and 64 diffusion weighted images at b = 700 s/mm^2^; 256 × 256 × 150 mm FOV and 2 mm isometric voxel size.

### Diffusion MRI processing

The diffusion weighted images were motion-corrected using FSL eddy (Andersson and Sotiropoulos, 2016) and upsampled to an isotropic voxel size of 1 mm using cubic spline interpolation. A diffusion tensor was fitted to the data of each voxel using weighted linear least squares (Veraart et al., 2013) and the results used to compute the fractional anisotropy map. The fiber orientation distribution functions were computed using constrained spherical deconvolution (Tournier et al., 2008) with a maximum spherical harmonics order of 8. The T1 weighted image was non-linearly registered to the b=0 image and fractional anisotropy map simultaneously using Advanced Normalization Tools (ANTs) (Tustison et al., 2014). The transformed T1 image was segmented using FreeSurfer to identify the cortical regions of the Desikan-Killiany atlas. The following cortical regions (See Suppl. Figure 1) were identified as target ROIs for streamlines: superior frontal (SF), caudal anterior cingulate (CAC), and rostral middle frontal (RMF). These ROIs were targeted since the literature showed that WM projections between the thalamus and the frontal cortex were particularly associated with frontal spindling activity (Gaudreault et al., 2018; Piantoni et al., 2013). Because of its size, the SF region was divided into three subregions of equal length in the anterior-posterior direction (anterior [SFa], middle [SFm], and posterior [SFp]), for a total of five ROIs. We specifically targeted fronto-central regions to avoid identifying corticospinal projections that target the pre-central and post-central regions but do not relay through the thalamus. The thalamus, whose segmentation was also provided by FreeSurfer, was identified as the starting region of interest. The FreeSurfer segmentation was also used to generate the tracking domain used in anatomically constrained particle filter tractography (Girard et al., 2014). More precisely, the “include” map, which dictates the brain regions where valid streamlines can end, was defined as the target ROIs. The “exclude” map, which dictates where streamlines cannot terminate, was defined as the complement of the “include” map, the cerebral WM of both hemispheres, the corpus callosum and the target ROIs. Finally, probabilistic tractography was initiated in the starting region using 10 seeds per voxel, a step size of 0.2 mm, and a maximum angle of 20 degrees between steps. Briefly, the tracking strategy above identifies streamlines that begin in the start region, end in the target region, and remain in the WM throughout their length. This procedure yielded 10 streamline bundles representing WM bundles starting in the thalamus and ending in one of the ROIs. For each bundle, the median length was used for the analyses since this measure is less influenced by outliers.

### Statistical analyses

First, ANOVAs with one independent factor (Sex: women vs. men) and one repeated measure (Derivations: F3, F4, C3, and C4) were performed. The same statistical analyzes were used to assess the interaction between sex and ROIs on the median length of the WM fiber bundles in millimeters (Sex: women vs. men; ROIs: CAC, RMF, SFa, SFm, and SFp). These analyzes were carried out independently for both hemispheres. Principal effects were further analyzed using post-hoc Tukey tests. The significance threshold was p < 0.05. As for all subsequent analyzes, we used the inverse of the median length in millimeters (i.e., 1/length) in our statistical models. As mentioned in the introduction, this choice is based on our objective to assess the propagation model and the anatomo-functional relationship between SS frequency and the length of the WM projections. The frequency of SS associated with a traveling wave over the thalamo-cortical loop must be related to the wavelength (*f · λ* = *v*) that corresponds to twice the length (L) of the WM projections (thalamo-cortical and cortico-thalamic projections) (*f ·* 2*L* = *v*). Therefore, the frequency of oscillation should regress with the inverse of the length between the thalamus and the cortex 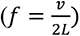.

#### Preliminary regression analyzes

We first carried out preliminary linear regressions between lengths of WM fiber bundles of interest and SS characteristics (frequency and amplitude) for each electrode (F3, F4, C3, C4) (See Supplementary materials for results). To account for multiple comparisons (2 hemispheres × 4 electrodes × 5 ROIs × 2 SS characteristics = 80 comparisons), we applied a Bonferroni correction and considered significance at p < 0.000625 (0.05 / 80 = 0.000625). Fourteen comparisons reached corrected significance (see Supplementary Materials and Suppl. Table 1 for complete results): significant associations were found between the left RMF and SS frequency on F4, the left SFa and SS frequency on F3, F4, C3, and C4, the left SFm and SS frequency on F3, F4, and C4, the right SFa and SS frequency on F4, C3, and C4, and finally the right SFm and SS frequency on F4, C3, and C4. Interestingly, no significant associations were observed between length of bundles and SS amplitude.

**Table 1.** Sex and derivation effects for sleep spindle frequency and amplitude in women and men

|  | Women<br>N = 14 | Men<br>N = 16 | Main effects (p values) |  |  | Effects |
| --- | --- | --- | --- | --- | --- | --- |
|  |  |  | Sex | Derivation | Interaction |  |
| <u>Spindle frequency (Hz)</u> |  |  | < 0.001 | < 0.001 | n.s. | W > M** |
| F3 | 12.9 ± 0.4 | 12.4 ± 0.2 |  |  |  | F3 < C3** |
| F4 | 12.8 ± 0.3 | 12.4 ± 0.2 |  |  |  | F3 < C4** |
| C3 | 13.2 ± 0.3 | 12.9 ± 0.4 |  |  |  | F4 < C3** |
| C4 | 13.3 ± 0.2 | 12.8 ± 0.3 |  |  |  | F4 < C4** |
| <u>Spindle amplitude (μV)</u> |  |  | < 0.001 | n.s. | n.s. | M < W** |
| F3 | 46.2 ± 13.1 | 35.1 ± 7.9 |  |  |  |  |
| F4 | 48.4 ± 16.6 | 35.1 ± 8.0 |  |  |  |  |
| C3 | 38.1 ± 8.9 | 34.3 ± 8.8 |  |  |  |  |
| C4 | 39.6 ± 10.8 | 32.7 ± 8.5 |  |  |  |  |

**Notes:** Data expressed as mean ± SD. P values were considered significant at $p < 0.05$ . n.s., non-significant.
One outlying value was removed from the C3 spindle amplitude analysis. \* = $p < 0.05$ , \*\* = $p < 0.01$

#### Main regression analyzes

To reduce the number of regression models, we performed subsequent analyzes using only the 14 pairs of variables (length of bundles/spindle frequency) showing significant associations in our conservatively corrected preliminary analyses. A second set of regression analyzes was performed to assess whether the median length of the WM fiber bundles between the thalamus and the frontal cortex predict SS variables when we controlled for the effects of sex and intracranial volume (ICV). More precisely, hierarchical linear regressions were used while entering sex and ICV as controlled variables in steps 1 and 2 respectively, and the median length of the WM fiber bundles in step 3. These covariates were added to the statistical model since significant sex-related differences were observed in our sample for both SS frequency and WM fiber lengths in addition to having been previously reported in the literature (Carrier et al., 2001; Martin et al., 2013; Menzler et al., 2011; Takao et al., 2014; Ujma et al., 2014). However, even if controlling for ICV is crucial in brain analyzes especially when sex is involved, it remains unclear whether statistical redundancy is present while correcting for both sex and ICV. Therefore, these covariates were added consecutively to the hierarchical statistical model to measure the additional explained variance of the ICV. This second covariate was also added to the model to ensure that the significant effects observed cannot rely only on head size but can rather be explained by the actual interindividual differences in the WM fiber lengths. Level of significance was set at p < 0.05. Moreover, we statistically tested the presence of multicollinearity between our predictors (i.e., the inverse of the median length of the WM fiber bundles, sex, and ICV) using the tolerance coefficient threshold of 0.1 and the variance inflation factor (VIF) threshold of 10.

Finally, a third set of analyzes aimed at investigating the effect of sex on SS frequency through the sex-related differences in the length of the WM fiber bundles underlying the thalamo-cortical loop. In other terms, it aimed at investigating the mediating effect of the length of the WM fiber bundles between the thalamus and the frontal cortex on the relationship between sex and SS frequency. These mediation analyzes thus tested the indirect effect of sex on SS frequency through its known direct effect on the length of the WM fiber bundles. The mediation analyzes were carried out with the SPSS macro PROCESS (v3.1) using 5,000 stratified bootstrap resamples to determine the bias-corrected 95^th^ percentile confidence intervals (Hayes, 2018). The mediations were based on the framework of model 4 as reported by Hayes (2018) for the computation of parallel mediation (See Suppl. Figure 2). First, sex was added to the model as a categorical variable. ICV and the length of the WM fiber bundles were then added to the model as separate parallel mediators even if our hierarchical regressions showed that ICV did not significantly explain more variance of SS frequency than did sex except for one regression between SS frequency on F4 and the left SFa. This model allows the computation of two specific indirect effects, namely an indirect effect of sex on SS frequency via the length of the WM fiber bundles only (a_1_·b_1_) and via the ICV only (a_2_·b_2_). A significance threshold of p < 0.05 was used in the interpretation of the total (c) and direct (c’) effects and a 95% bias-corrected confidence interval not including zero was used to interpret significant mediation of indirect effects (Hayes, 2018).

**Figure 2.**
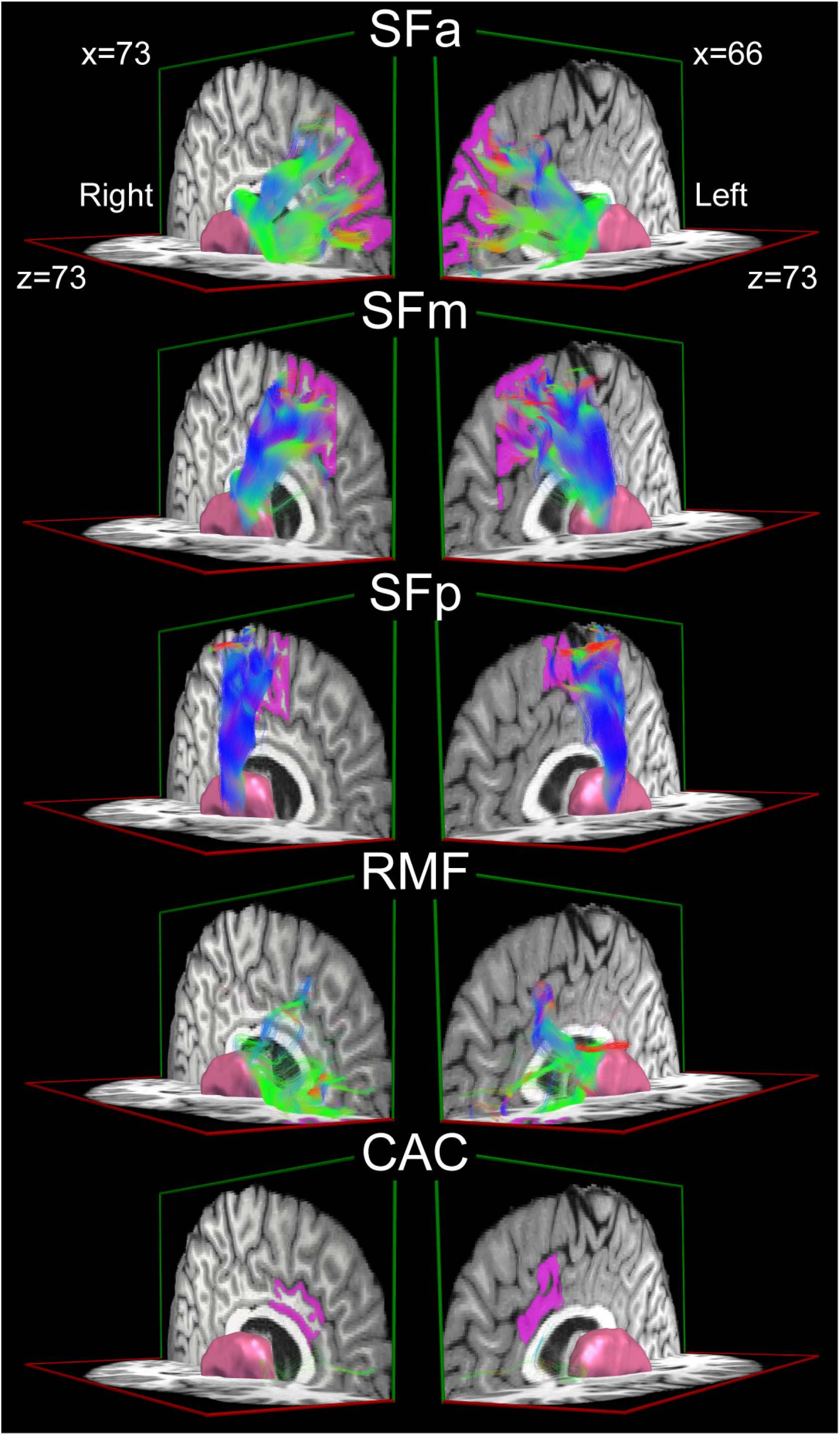
White matter fiber bundles connecting the thalamus to different regions of interest in the frontal cortex. Representation of the white matter (WM) fiber bundles connecting the thalamus (in pink) to our pre-selected regions of interest (ROIs; in magenta) derived from our tractography pipeline for one particular subject. The streamlines are overlaid on the individual’s preprocessed T1w image and color-encoded according to the fiber orientation map. Fibers predominantly oriented in the left-right axis are in red, in the anterior-posterior axis in green, and in the superior-inferior axis in blue. (SFa: anterior part of the superior frontal gyrus, SFm: middle part of the superior frontal gyrus, SFp: posterior part of the superior frontal gyrus, RMF: rostral middle frontal gyrus, CAC: caudal anterior cingulate gyrus)

## Results

### Sex differences on sleep spindle variables and length of white matter fiber bundles

Sex and derivation effects on SS frequency and amplitude are presented in Table 1. Women showed higher SS frequency compared to men (Sex effect: F(1,112) = 69.8, p < 0.001). SS frequency was significantly higher in central derivations compared to frontal electrodes (Derivation effect: F(3,112) = 23.8, p < 0.001). No significant interaction between sex and derivation was found for SS frequency. Women also showed higher amplitude than men (F(1,111) = 20.3, p < 0.001) but no significant effect of derivation was observed for spindle amplitude.. Significant sex effects were also found for the median length of WM fiber bundles where men showed significantly longer WM streamlines compared to women in each hemisphere (Left hemisphere: F(1,128) = 8.2, p < 0.01; Right hemisphere: F(1,130) = 17.0, p < 0.001; see Table 2). Significant ROIs effects were also found (Left hemisphere: F(4,128) = 11.5, p < 0.001; Right hemisphere: F(4,130) = 15.7, p < 0.001). Globally, the tracts between the thalamus and the CAC were the longest whereas the tracts to the SFp were the shortest bundles. No significant interaction between sex and target ROIs was found for WM median fiber bundle length.

**Table 2.** Median length of white matter fiber bundles from the thalamus to the frontal cortex in women and men.

|  | Women<br>N = 14 | Men<br>N = 16 | Main effects (p values) |  |  | Effects |
| --- | --- | --- | --- | --- | --- | --- |
|  |  |  | Sex | ROIs | Interaction |  |
| <u>Left hemisphere ROIs</u> |  |  | < 0.01 | < 0.001 | n.s. | W < M** |
| CAC | 95.2 ± 41.1 | 106.5 ± 38.0 |  |  |  | CAC > RMF** |
| RMF | 80.6 ± 6.0 | 86.5 ± 3.8 |  |  |  | CAC > SFa* |
| SFa | 83.3 ± 4.8 | 92.4 ± 4.7 |  |  |  | CAC > SFm** |
| SFm | 73.5 ± 3.9 | 81.7 ± 3.6 |  |  |  | CAC > SFp** |
| SFp | 69.2 ± 2.9 | 75.0 ± 3.7 |  |  |  | SFa > SFp** |
| <u>Right hemisphere ROIs</u> |  |  | < 0.001 | < 0.001 | n.s. | W < M** |
| CAC | 83.9 ± 21.2 | 95.3 ± 24.8 |  |  |  | CAC > SFm** |
| RMF | 82.0 ± 4.2 | 86.7 ± 5.1 |  |  |  | CAC > SFp** |
| SFa | 84.1 ± 4.5 | 92.1 ± 4.5 |  |  |  | RMF > SFp** |
| SFm | 73.4 ± 3.7 | 81.4 ± 4.9 |  |  |  | SFa > SFm** |
| SFp | 68.2 ± 3.0 | 73.4 ± 3.7 |  |  |  | SFa > SFp** |

**Notes:** Data expressed as mean ± SD. The median length is expressed in millimeters (mm). P values were considered significant at $p < 0.05$ . CAC, caudal anterior cingulate, n.s., non-significant, ROIs, regions of interests, RMF, rostral middle frontal, SFa, superior frontal anterior, SFm, superior frontal middle, SFp, superior frontal posterior. \* = $p < 0.05$ , \*\* = $p < 0.01$

### Preliminary linear regressions

Preliminary linear regressions (Suppl. Table 1) were carried out on the WM fiber bundles of interest and sleep spindles (SS) characteristics (frequency and amplitude) on each electrode (F3, F4, C3, C4). Fourteen comparisons reached the Bonferroni correction of p < 0.000625: the left RMF predicted SS frequency on F4 (R^2^ = 0.498, β = 0.706, p = 0.000081), the left SFa predicted SS frequency on F3 (R^2^ = 0.400, β = 0.632, p = 0.000179), F4 (R^2^ = 0.569, β = 0.754, p = 0.000001), C3 (R^2^ = 0.461, β = 0.679, p = 0.000037), and C4 (R^2^ = 0.504, β = 0.710, p = 0.000011), the left SFm predicted SS frequency on F3 (R^2^ = 0.465, β = 0.682, p = 0.000033), F4 (R^2^ = 0.517, β = 0.719, p = 0.000008), and C4 (R^2^ = 0.476, β = 0.690, p = 0.000025), the right SFa predicted SS frequency on F4 (R^2^ = 0.410, β = 0.640, p = 0.000138), C3 (R^2^ = 0.415, β = 0.645, p = 0.000121), and C4 (R^2^ = 0.476, β = 0.629, p = 0.000195), and the right SFm predicted SS frequency on F4 (R^2^ = 0.388, β = 0.623, p = 0.000239), C3 (R^2^ = 0.350, β = 0.592, p = 0.000573), and C4 (R^2^ = 0.503, β = 0.709, p = 0.000011).

### The inverse of the length (1/L) of white matter fiber bundles predicts sleep spindle frequency

Significant associations between the inverse of the median length of the WM fiber bundles and SS frequency were found when adding sex and ICV as consecutive control variables in hierarchical regression models (See Tables 3 and 4). Figure 2 shows the fiber bundles identified with tractography from the thalamus to each ROIs for one specific individual in our study. Globally, longer WM bundles between the thalamus and the frontal lobe were associated with lower frontal SS frequency. More precisely, the inverse of the length of the WM fiber bundles between the thalamus and both the left SFa and left RMF were positively correlated with SS frequency in F4 derivations (Figure 3A; SFa: β = 0.465, p = 0.034; RMF: β = 0. 482, p = 0.020) and explained respectively 7.5 and 12.8% of the variance. The inverse of the median length of WM bundles connecting the thalamus to the left and right SFa regions was positively correlated with C3 SS frequency (Figure 3B; Left SFa: β = 0.638, p = 0.014; Right SFa: β = 0. 496, p = 0.030) whereas the projections to the left SFa, and the right SFm were positively correlated with SS frequency for the C4 electrode (Figure 3C; Left SFa: β = 0. 477, p = 0.039; Right SFm: β = 0. 542, p = 0.032) explaining between 7.1 and 14.1% of the variance. None of the hierarchical regressions carried out on F3 reached significance. Also, the median length of the WM fiber bundles was not significantly associated with SS amplitude.

**Table 3.** Hierarchical regression analyses with the inverse of the median length of white matter fiber bundles from the thalamus to the frontal cortex as predictor of sleep spindle frequency on the F4 electrode.

| Predictors | <u>F4 sleep spindle frequency</u> |  |  |
| --- | --- | --- | --- |
| | $\beta$ | $R^2$ | $R^2$ change |
| *Step 1* |  | 34.5%** |  |
| Sex | 0.587** |  |  |
| *Step 2* |  | 44.2%** | 9.7% |
| ICV | -0.444 |  |  |
| *Step 3* |  | 57.0%** | 12.8%* |
| Left RMF | 0.482* |  |  |
| Predictors | $\beta$ | $R^2$ | $R^2$ change |
| *Step 1* |  | 41.9%** |  |
| Sex | 0.648** |  |  |
| *Step 2* |  | 53.9%** | 11.9%* |
| ICV | -0.526* |  |  |
| *Step 3* |  | 61.3%** | 7.5%* |
| Left SFa | 0.465* |  |  |

**Notes:** This hierarchical regression model was carried out using the inverse of the median length in millimeters (i.e. 1/length). ICV, Intracranial volume; RMF, rostral middle frontal; SFa, anterior part of the superior frontal gyrus. \* $p < 0.05$ , \*\* $p < 0.01$

**Table 4.** Hierarchical regression analyses with the inverse of the median length of white matter fiber bundles from the thalamus to the frontal cortex as predictor of sleep spindle frequency on the C3 and C4 electrodes.

| Predictors | C3 sleep spindle frequency |  |  | C4 sleep spindle frequency |  |  |
| --- | --- | --- | --- | --- | --- | --- |
| | $\beta$ | $R^2$ | $R^2$ change | $\beta$ | $R^2$ | $R^2$ change |
| *Step 1* |  | 28.7%** |  |  | 45.7%** |  |
| Sex | 0.536** |  |  | 0.676** |  |  |
| *Step 2* |  | 32.8%** | 4.1% |  | 48.7%** | 3.0% |
| ICV | -0.308 |  |  | -0.264 |  |  |
| *Step 3* |  | 46.9%** | 14.1%* |  | 56.6%** | 7.9%* |
| Left SFa | 0.638* |  |  | 0.477* |  |  |
| Predictors | $\beta$ | $R^2$ | $R^2$ change | $\beta$ | $R^2$ | $R^2$ change |
| *Step 1* |  | 28.7%** |  | -- | -- | -- |
| Sex | 0.536** |  |  | -- | -- | -- |
| *Step 2* |  | 32.8%** | 4.1% | -- | -- | -- |
| ICV | -0.308 |  |  | -- | -- | -- |
| *Step 3* |  | 44.2%** | 11.4%* | -- | -- | -- |
| Right SFa | 0.496* |  |  |  |  |  |
| Predictors | $\beta$ | $R^2$ | $R^2$ change | $\beta$ | $R^2$ | $R^2$ change |
| *Step 1* | -- | -- | -- |  | 45.7%** |  |
| Sex |  |  |  | 0.676** |  |  |
| *Step 2* | -- | -- | -- |  | 48.7%** | 3.0% |
| ICV |  |  |  | -0.264 |  |  |
| *Step 3* | -- | -- | -- |  | 57.1%** | 8.5%* |
| Right SFm |  |  |  | 0.542* |  |  |

**Notes:** This hierarchical regression model was carried out using the inverse of the median length in millimeters (i.e. 1/length). ICV, Intracranial volume; SFa, anterior part of the superior frontal gyrus; SFm, middle part of the superior frontal gyrus. \* $p < 0.05$ , \*\* $p < 0.01$

**Figure 3.**
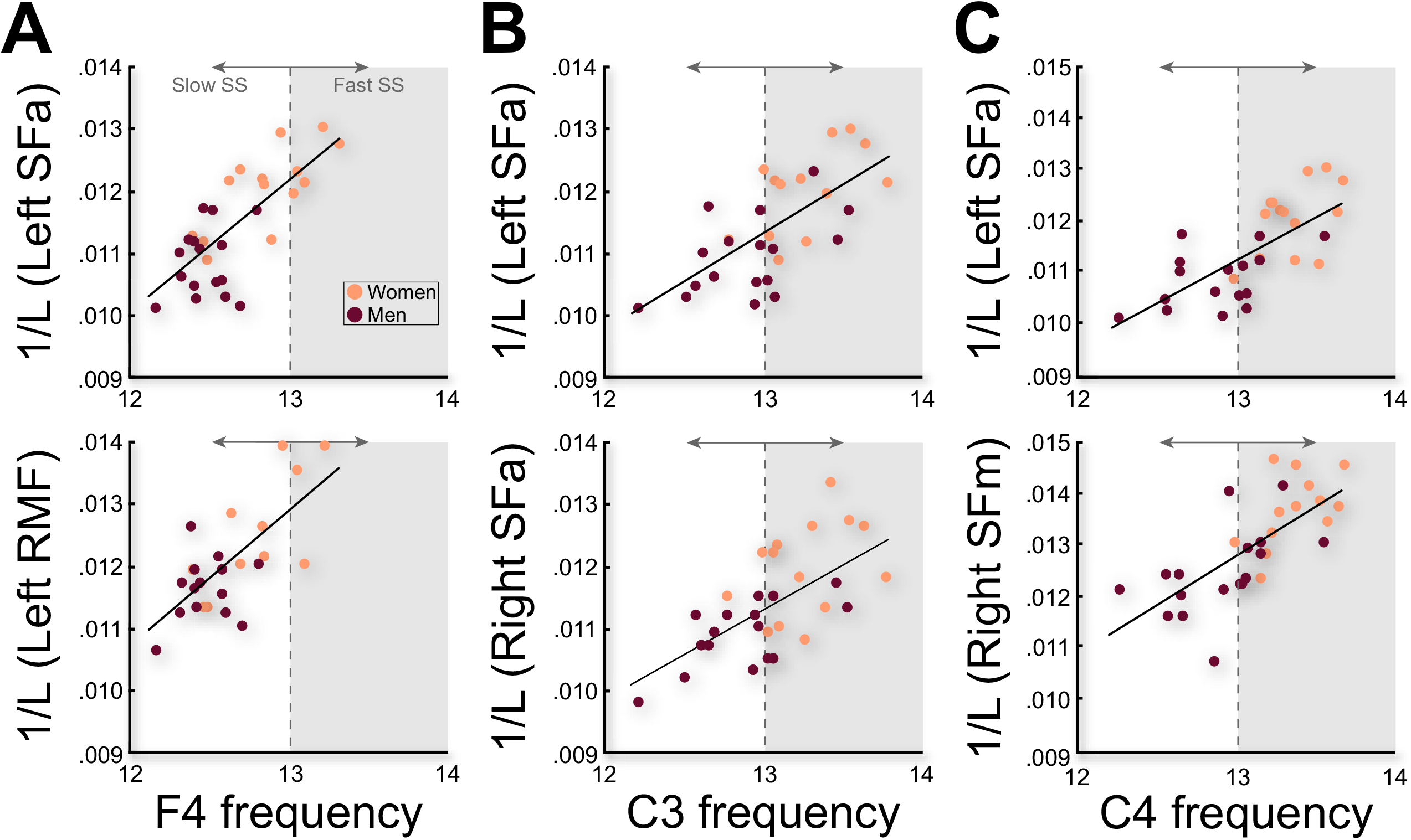
Scatterplots of the association between white matter (WM) fiber bundles and sleep spindle frequency (SS). Significant associations between the inverse of the median length of the WM fiber bundles and SS frequency respectively on F4 (A), C3 (B), and C4 (C) electrodes.

### Mediation effect of the inverse of the length (1/L) of the thalamo-cortical white matter fiber bundles on sex-related differences in sleep spindle frequency

In order to evaluate whether the inverse of the length of WM fiber bundles could predict sex-related differences in SS frequency, we performed mediation analyzes (results are presented in Table 5). The mediation analyzes showed a significant indirect effect of sex on F4 SS frequency through the mediating effect of the inverse of the length of WM fiber bundles from the thalamus to the left RMF (a_1_·b_1_ = 0.142, 95% CI = 0.014 – 0.332), and to the left SFa (a_1_·b_1_ = 0.195, 95% CI = 0.038 – 0.404). More specifically, shorter WM projections between the thalamus and the frontal cortex significantly predicted faster SS frequency on the F4 derivation in women as compared to men. We observed a similar indirect effect of sex on C3 SS frequency via the inverse of the length of the projections connecting the thalamus to the left (a_1_·b_1_ = 0.330, 95% CI = 0.066 – 0.677) and right SFa (a_1_·b_1_ = 0.233, 95% CI = 0.054 – 0.548) as well as on C4 SS frequency via the fiber bundles targeting the right SFm (a_1_·b_1_ = 0.269, 95% CI = 0.055 – 0.566). Moreover, all direct effects were nonsignificant in these mediation analyzes, suggesting that the inverse of the length of WM fiber bundles completely mediated the effect of sex on SS frequency. Taken together, these results suggest that sex-related differences observed in SS frequency on frontal and central derivations are predicted by the inverse of the length of WM fiber bundles connecting the thalamus to the left RMF, bilateral SFa, and right SFm. No significant indirect effect was found with the ICV indicating that the sex-related difference in SS frequency was not predicted through its effect on ICV.

**Table 5.** Mediation analyses for the relationships of sex on sleep spindle frequency via the inverse of the median length of the white matter fiber bundles connecting the thalamus to the frontal cortex and the intracranial volume

| Tracts | Laterality | Derivation | c | a<sub>1</sub> | b<sub>1</sub> | a<sub>2</sub> | b<sub>2</sub> | c' | Indirect effects (95% CI) |  |
| --- | --- | --- | --- | --- | --- | --- | --- | --- | --- | --- |
|  |  |  |  |  |  |  |  |  | WM (a<sub>1</sub>·b<sub>1</sub>) | ICV (a<sub>2</sub>·b<sub>2</sub>) |
| RMF | L | F4 | 0.322** | 0.0009** | 158.206* | -0.234*** | -0.283 | 0.114 | 0.142 (0.014–0.332)* | 0.066 (-0.132–0.215) |
| SFa | L | F4 | 0.385*** | 0.0012*** | 163.094* | -0.252*** | -0.435 | 0.081 | 0.195 (0.038–0.404)* | 0.110 (-0.085–0.297) |
| SFa | L | C3 | 0.393** | 0.0012*** | 276.108* | -0.252*** | -0.169 | 0.106 | 0.330 (0.066–0.677)* | -0.043 (-0.360–0.219) |
| SFa | R | C3 | 0.393** | 0.0010*** | 221.696* | -0.252*** | -0.082 | 0.140 | 0.233 (0.054–0.548)* | 0.021 (-0.404–0.219) |
| SFa | L | C4 | 0.482*** | 0.0012*** | 200.520* | -0.252*** | 0.051 | 0.255 | 0.239 (-0.006–0.567) | -0.013 (-0.273–0.228) |
| SFm | R | C4 | 0.482*** | 0.0013*** | 202.659* | -0.252*** | 0.275 | 0.282 | 0.269 (0.055–0.566) * | -0.070 (-0.370–0.212) |

**Notes:** c = total effect of sex on sleep spindle (SS) frequency; a<sub>1</sub> = effect of sex on median length of white matter tracts (WM, mediator 1); b<sub>1</sub> = association between WM tracts and SS frequency; a<sub>2</sub> = effect of sex on intracranial volume (ICV, mediator 2); b<sub>2</sub> = association between ICV and SS frequency; c' = direct effect of sex on SS frequency; RMF, Rostral middle frontal cortex; SFa, Anterior part of the superior frontal cortex; SFm, Middle part of the superior frontal cortex. For white matter tracts, the inverse of the median length in millimeters (i.e. 1/length) was used in the mediation analyses. CI = 95% bootstrap confidence interval for indirect effect (5,000 stratified resamples). \*p < 0.05 \*\*p < 0.01 \*\*\*p < 0.001

## Discussion

The neurophysiological underpinnings of inter-individual sleep EEG variability and more specifically of sex-related differences in SS characteristics are still poorly understood. The goal of our study was to test a propagation model of a signal along the WM fibers generating SS. We then assessed the relationship between the inverse of the length of the WM fiber bundles underlying the thalamo-cortical loop (an anatomical measure) and the SS frequency (which can be considered as a functional representation of this system). We used a precise streamline-based and bundle-specific approach to quantitatively measure the length of the WM fiber bundles between the thalamus and specific ROIs in the frontal cortex. Our main results showed that longer WM fiber bundles between the thalamus and the frontal lobe, more precisely the left RMF, the bilateral SFa, and the right SFm were associated with lower SS frequencies in frontal and central parasagittal derivations. Moreover, our mediation analyzes showed that the sex-related differences observed in SS frequency were completely mediated by the length of the WM projections from the thalamus to the frontal cortex. To our knowledge, this is the first time a neuroanatomical marker is associated with inter-individual variability in SS frequency and may explain sex-related differences in SS frequency.

### Thalamo-cortical projection length predicts sleep spindle frequency

Anatomically, the distance separating the thalamus from the cerebral cortex is thought to be analog to the path taken by the nerve impulses back and forth along the thalamo-cortical loop. Assuming that SS are network-related events, all nuclei (i.e., the neuronal relays) implicated in this loop must be finely tuned for their waxing and waning properties. From the burst activity originating from the neurons of the thalamic reticular nucleus to the cortical cells, multiple processes are thereafter initiated, including the activation of I_h_ (hyperpolarization-activated cation currents) which causes a rapid depolarization of the membrane, leading to bursts of excitatory action potentials in thalamo-cortical neurons (McCormick and Bal, 1997). The action potentials that reach the cortex via these projections increase the probability of depolarizing the cortico-thalamic neurons which, in turn, project back to the thalamus keeping the spindling oscillations. However, cortico-thalamic projections also cause a gradual depolarization of the membrane of thalamo-cortical neurons which, over time, prevents the activation of I_h_ currents and leads to a disruption of the rhythmic activity in the loop causing the SS waning and the inter-SS refractory period (Bal and McCormick, 1996; Lüthi and McCormick, 1998). Therefore, we consider SS as being a cortical response of a synchronized activity stimulated by the local oscillation of a traveling wave propagating along the thalamo-cortical loop; this model should relate shorter thalamic projections to the cortex with faster back and forth in the network, before the desynchronization of the cortical process. Conversely, longer WM fibers between the thalamus and the cortex would involve more time to complete the reciprocal action potentials in the loop, which would limit the number of cycles in the time needed for the whole system to fade and terminate the SS. Our study therefore brings supporting evidence that SS frequency recorded on the scalp (a functional substrate of this well-described system) is indeed representative of the propagation of nerve impulses on the anatomical projections between the thalamus and the cortex. Moreover, based on the proposed model (*v*/*f* = *λ* where *λ* = 2*L*), nerve impulse speed can be estimated from the WM fiber bundle lengths and the SS frequency measured in our sample. The estimated velocities are ranging from 1.5 to 2.7 m/s which can be considered of plausible magnitude according to the literature (Deslauriers-Gauthier and Deriche, 2019; Fukushima et al., 2015; Innocenti et al., 2015). It is however of note that our model was fairly simple and did not consider any integration or conduction delays, which could lead to a more accurate modeling of the nerve impulse speed (Caminiti et al., 2013; Innocenti et al., 2014). Although further methodological development in the field is needed at the moment to estimate such parameters in vivo, it is likely that these other factors affecting nerve impulse conduction (i.e., integration delay, myelin integrity, internodal spacing, and axon diameter) may contribute to the variance of SS frequency, and so, even in healthy WM projections (Horowitz et al., 2015). For instance, a recent probabilistic tractography study characterized fiber length distribution across the human brain and found the density of fibers of specific length to be different depending on the brain area, but they also showed greater myelin content in shorter fibers (Bajada et al., 2019). Most importantly, these factors may explain the presence of similar frequencies among brain oscillations in both humans and other species despite different brain sizes (Wang et al., 2008). More specifically, studies assessing axon diameters and conduction velocity/delays investigated the differences across species and showed that, even if both larger and smaller brains have a similar calibre of axon diameters, there is a specific increase of axon diameters in fibers associated with faster information processing among mammals with larger brain volume (Buzsáki et al., 2013; Caminiti et al., 2013; Liewald et al., 2014). Such neurophysiological and metabolic differences might contribute to the functional preservation of brain oscillations (Wang et al., 2008) and might as well reflect evolutionary processes (Caminiti et al., 2009).

Our results are also reflective of a novel and specific measure of diffusion MRI for assessing the physical length of thalamo-cortical WM projections. To ensure the independence between the length of the WM fiber bundles connecting the thalamus to the frontal cortex and the size of the head, we corrected for the total intracranial volume in all our statistical analyzes. Indeed, our results demonstrated that, while taking into account head size, the inverse of the length of the WM fiber bundles significantly explained more variance than only sex and/or ICV. This result was also corroborated by our mediation analysis showing no mediating effect of the ICV on sex-related changes in SS frequency whereas the inverse of the length of WM projections completely mediated the effect of sex on SS frequency. Finally, it is important to point out that our analyzes were carried out with the inverse of the median length (1/L) and not only the length between targets, and that the results of the regressions predicting SS frequency were slightly more significant using the inverse of L instead of L, which adds confidence that our results are not simply attributable to head size.

Our findings point to a neurophysiological mechanism that could predict the topographical differences in SS frequency over the scalp. Our data, showing a lower mean frequency in frontal than central derivations, corroborate the accepted notion that SS frequency increases according to an antero-posterior gradient (Cox et al., 2017; Dehghani et al., 2011; Martin et al., 2013). It is then possible to argue that SS in more posterior derivations are faster because connections between the thalamus and more posterior brain areas are shorter. Further studies are needed to investigate how projections from thalamus to other cortical ROIs may predict higher posterior SS frequency. For instance, whole-brain tractograms could bring more insight into the topographical modulation of SS frequency as well as into the specific implication of both ipsilateral and contralateral WM fiber bundles. In this regard, our study considered WM fiber bundles to be symmetrical between hemispheres which would increase the chances of finding contralateral correlations as well as the expected ipsilateral ones. The very conservative Bonferroni threshold used to empirically identify significant associations during the preliminary analyzes also decreased the possibility to assess the specificity of associations with only a subset of correlations reached the severe significance threshold. Such studies should also include complementary methods such as high-density EEG which would take into consideration co-occurring SS-related activity on multiple electrodes and would allow the modeling of volume conduction which our EEG data was susceptible to. Finally, these advanced methodologies would allow the assessment of the specificity and the validity of the thalamo-cortical projections (Maier-Hein et al., 2017). It is also interesting to note that other EEG waves such as alpha oscillations are known to be related to the thalamo-cortical loop as well and have been associated with the resonance properties of this system in computational models (Hindriks and van Putten, 2013). Further studies should therefore investigate whether WM characteristics also play a role in the frequency of these oscillations.

### Sex differences in sleep spindle frequency are mediated by the thalamo-cortical projections

The length of the WM fibers not only predicted SS frequency, but also completely mediated the sex-related differences in SS frequency. In other words, after adding the length of the WM fiber bundles to the model, the relationship between sex and SS frequency was no longer significant. To our knowledge, this is the first study demonstrating that neuroanatomical differences predict the sex-related SS frequency variation. Future studies should however include more women for replication purposes and aim at tracking their hormonal/contraceptive status as one limitation of our study is the recruitment of women participants regardless of their status. Although very few hypotheses have been stated to explain sex-related differences in sleep, sex steroids have been at the forefront as a possible mechanism since objective sleep measures and SS vary across the menstrual cycle and during pregnancy in women (Carrier et al., 2017; Sharkey et al., 2014). A significant impact of the premenstrual/menstrual phase in women has also been associated with subjective sleep quality, but also with the EEG power density in the SS-related sigma frequency band (Driver et al., 1996; Romans et al., 2015).

The length of the WM fiber bundles connecting the thalamus to the frontal cortex significantly predicted the frequency of SS recorded in frontal and central parasagittal electrodes, bringing a first quantitative measure of the involvement of the thalamo-cortical loop in the interindividual variation of SS frequency. Moreover, these specific thalamo-cortical projections also completely mediated the sex-related difference in SS frequency. In other words, these results suggest that the sexual dimorphism observed in SS frequency is statistically explained by the between-sex difference in length of the thalamo-cortical projections. Further studies will be needed to completely understand the implication of such a process in specific brain functions associated with SS such as sleep-dependent cognitive functions or even interindividual differences in fluid intelligence. Integrative models should be considered to investigate the implication of physical quantitative measures (i.e., length of the WM fiber bundles) as well as dMRI-derived WM metrics along these fibers on the modulation of SS variables.

## Acknowledgments

This study was supported by the Canadian Institutes of Health Research (CIHR – grant number 190750), the Natural Sciences and Engineering Council of Canada (NSERC – grant number RGPIN-2016-05149), and the Fonds de recherche du Québec – Santé (FRQS). The authors would like to thank Dominique Petit, PhD and Carrie Schipper for reviewing the manuscript.

## Data Availability

Data Availability: All codes and softwares used to analyze data are freely available and referenced in the manuscript. De-identified and transformed data used to produce all the figures in the manuscript are available upon request to the first (Pierre-Olivier Gaudreault) and corresponding author (Julie Carrier). The dataset cannot be shared as participants did not give consent for data sharing. To get access to the raw data, a request must be formulated to the ethics committee of the Hôpital de Sacré-Coeur de Montréal as raw data from human participants cannot be made public under Québec’s law. The data provided will be anonymized. Researchers who request access to the data will need to provide their research protocol and their IRB approval for their protocol. All documents will be studied by the owner of the database (Julie Carrier) who will also submit to the institution’s REB for authorization to share the data. Requests should be addressed to: Julie Carrier (PI): and Sonia Frenette (in cc).

## Author Contributions

P-O.G. contributed to the design of the analyses, statistical analyses, and writing of the manuscript; J-M.L. contributed to the design of the analyses and writing of the manuscript; M.D. contributed to the design of the analyses and the review of the manuscript; N.G. contributed to design of the study and the review of the manuscript; J.D. contributed to the review of the manuscript; S.D.G. contributed to the design of the analyses, to the data processing, and the writing and the review of the manuscript; J.C. contributed to the design of the study, the writing and the review of the manuscript.

## Figure Captions

**Suppl. Figure 1.**
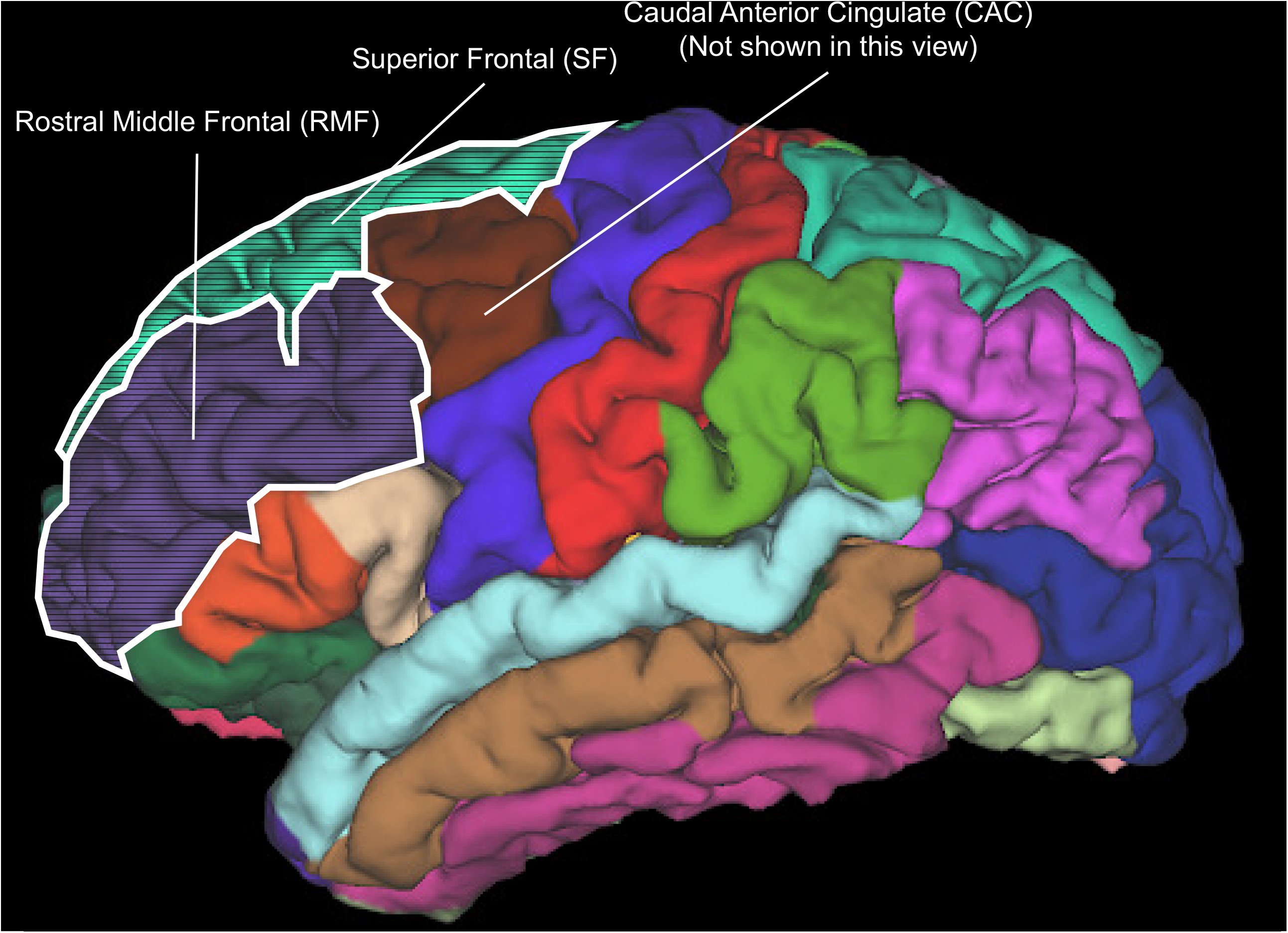
Representation of the regions of interest (ROIs). This figure represents the cortical ROIs targeted in this study overlaid on a standard brain template (ICBM152) parcellated using the Desikan-Killiany atlas (Desikan et al., 2006). The Caudal anterior cingulate (CAC) gyrus is situated inside the brain and is therefore not pictured on this figure. The Superior frontal (SF) region was further subdivided into the anterior (SFa), middle (SFm), and posterior parts (SFp).

**Suppl. Figure 2.**
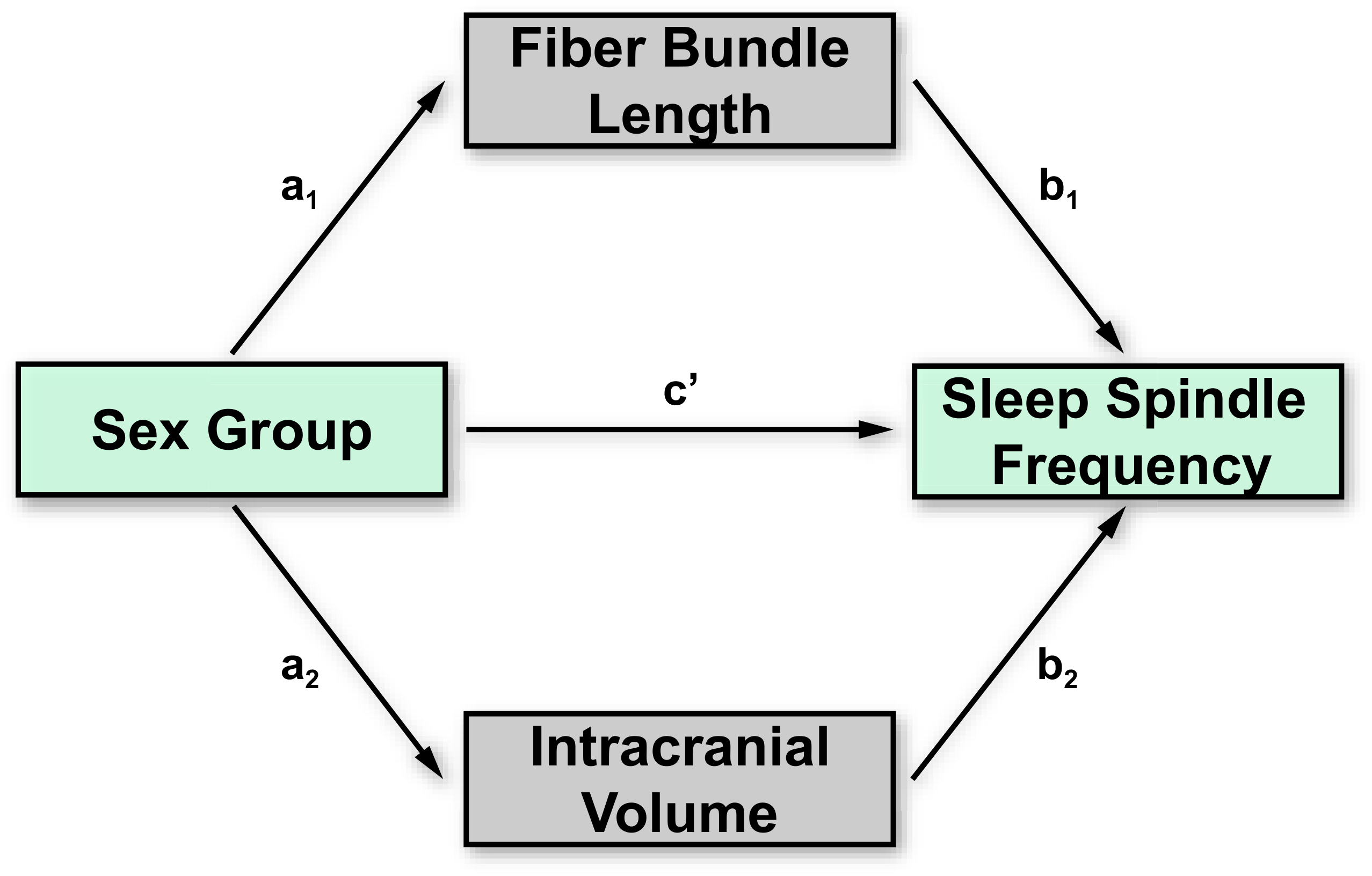
Mediation analysis model. Parallel mediation model used to investigate the indirect effect of sex on SS frequency through the fiber bundle length and the ICV. The ICV was added in the model as a control for brain size. This model allows the computation of two specific indirect effects, namely an indirect effect of sex on SS frequency via the length of the WM fiber bundles only (a_1_·b_1_) and via the ICV only (a_2_·b_2_).

## References

Andersson, J.L.R., Sotiropoulos, S.N. An integrated approach to correction for off-resonance effects and subject movement in diffusion MR imaging. NeuroImage, 2016, 125: 1063–1078.

Bajada, C.J., Schreiber, J., Caspers, S. Fiber length profiling: A novel approach to structural brain organization. NeuroImage, 2019, 186: 164–173.

Bal, T., McCormick, D.A. What Stops Synchronized Thalamocortical Oscillations? Neuron, 1996, 17: 297–308.

Blume, C., del Giudice, R., Wislowska, M., Heib, D.P.J., Schabus, M. Standing sentinel during human sleep: Continued evaluation of environmental stimuli in the absence of consciousness. NeuroImage, 2018, 178: 638–648.

Bonjean, M., Baker, T., Lemieux, M., Timofeev, I., Sejnowski, T., Bazhenov, M. Corticothalamic Feedback Controls Sleep Spindle Duration In Vivo. Journal of Neuroscience, 2011, 31: 9124–9134.

Buzsáki, G., Logothetis, N., Singer, W. Scaling Brain Size, Keeping Timing: Evolutionary Preservation of Brain Rhythms. Neuron, 2013, 80: 751–764.

Caminiti, R., Carducci, F., Piervincenzi, C., et al. Diameter, Length, Speed, and Conduction Delay of Callosal Axons in Macaque Monkeys and Humans: Comparing Data from Histology and Magnetic Resonance Imaging Diffusion Tractography. Journal of Neuroscience, 2013, 33: 14501–14511.

Caminiti, R., Ghaziri, H., Galuske, R., Hof, P.R., Innocenti, G.M. Evolution amplified processing with temporally dispersed slow neuronal connectivity in primates. Proc. Natl. Acad. Sci. U.S.A., 2009, 106: 19551–19556.

Carrier, J., Land, S., Buysse, D.J., Kupfer, D.J., Monk, T.H. The effects of age and gender on sleep EEG power spectral density in the middle years of life (ages 20-60 years old). Psychophysiology, 2001, 38: 232–242.

Carrier, J., Semba, K., Deurveilher, S., et al. Sex differences in age-related changes in the sleep-wake cycle. Frontiers in Neuroendocrinology, 2017, 47: 66–85.

Clawson, B.C., Durkin, J., Aton, S.J. Form and Function of Sleep Spindles across the Lifespan. Neural Plasticity, 2016, 2016: 1–16.

Cox, R., Schapiro, A.C., Manoach, D.S., Stickgold, R. Individual Differences in Frequency and Topography of Slow and Fast Sleep Spindles. Front. Hum. Neurosci., 2017, 11: 433.

De Gennaro, L., Ferrara, M. Sleep spindles: an overview. Sleep Medicine Reviews, 2003, 7: 423–440.

Dehghani, N., Cash, S.S., Halgren, E. Topographical frequency dynamics within EEG and MEG sleep spindles. Clinical Neurophysiology, 2011, 122: 229–235.

Deslauriers-Gauthier, S., Deriche, R. Estimation of axonal conduction speed and the inter hemispheric transfer time using connectivity informed maximum entropy on the mean. In: Gimi, B. and Krol, A. (eds.) Medical Imaging 2019: Biomedical Applications in Molecular, Structural, and Functional Imaging. p. 12. SPIE, San Diego, United States (2019).

Drakesmith, M., Harms, R., Rudrapatna, S.U., Parker, G.D., Evans, C.J., Jones, D.K. Estimating axon conduction velocity in vivo from microstructural MRI. NeuroImage, 2019, 203: 116186.

Driver, H.S., Dijk, D.J., Werth, E., Biedermann, K., Borbély, A.A. Sleep and the sleep electroencephalogram across the menstrual cycle in young healthy women. Journal of Clinical Endocrinology and Metabolism, 1996, 81: 728–735.

Fang, Z., Sergeeva, V., Ray, L.B., Viczko, J., Owen, A.M., Fogel, S.M. Sleep Spindles and Intellectual Ability: Epiphenomenon or Directly Related? Journal of Cognitive Neuroscience, 2017, 29: 167–182.

Fernandez, L.M.J., Lüthi, A. Sleep Spindles: Mechanisms and Functions. Physiological Reviews, 2020, 100: 805–868.

Fogel, S.M., Smith, C.T. The function of the sleep spindle: A physiological index of intelligence and a mechanism for sleep-dependent memory consolidation. Neuroscience & Biobehavioral Reviews, 2011, 35: 1154–1165.

Fukushima, M., Yamashita, O., Knösche, T.R., Sato, M. MEG source reconstruction based on identification of directed source interactions on whole-brain anatomical networks. NeuroImage, 2015, 105: 408–427.

Gaudreault, P.-O., Gosselin, N., Lafortune, M., et al. The association between white matter and sleep spindles differs in young and older individuals. Sleep, 2018, 41:.

Girard, G., Whittingstall, K., Deriche, R., Descoteaux, M. Towards quantitative connectivity analysis: reducing tractography biases. NeuroImage, 2014, 98: 266–278.

Hayes, A.F. Introduction to mediation, moderation, and conditional process analysis: a regression-based approach. Guilford Press, New York, 2018.

Hindriks, R., van Putten, M.J.A.M. Thalamo-cortical mechanisms underlying changes in amplitude and frequency of human alpha oscillations. NeuroImage, 2013, 70: 150–163.

Horowitz, A., Barazany, D., Tavor, I., Bernstein, M., Yovel, G., Assaf, Y. In vivo correlation between axon diameter and conduction velocity in the human brain. Brain Struct Funct, 2015, 220: 1777–1788.

Iber, C., Ancoli-Israel, S., Chesson, A.L., Quan, S.F. The AASM Manual for the Scoring of Sleep and Associated Events: Rules, Terminology, and Technical Specifications. American Academy of Sleep Medicine, Westchester, IL, 2007.

Innocenti, G.M., Caminiti, R., Aboitiz, F. Comments on the paper by Horowitz et al. (2014). Brain Struct Funct, 2015, 220: 1789–1790.

Innocenti, G.M., Vercelli, A., Caminiti, R. The Diameter of Cortical Axons Depends Both on the Area of Origin and Target. Cerebral Cortex, 2014, 24: 2178–2188.

Kahana, M.J. The Cognitive Correlates of Human Brain Oscillations. J. Neurosci., 2006, 26: 1669–1672.

Liewald, D., Miller, R., Logothetis, N., Wagner, H.-J., Schüz, A. Distribution of axon diameters in cortical white matter: an electron-microscopic study on three human brains and a macaque. Biol Cybern, 2014, 108: 541–557.

Lüthi, A. Sleep Spindles: Where They Come From, What They Do. Neuroscientist, 2014, 20: 243–256.

Lüthi, A., McCormick, D.A. Periodicity of Thalamic Synchronized Oscillations: the Role of Ca2+-Mediated Upregulation of Ih. Neuron, 1998, 20: 553–563.

Maier-Hein, K.H., Neher, P.F., Houde, J.-C., et al. The challenge of mapping the human connectome based on diffusion tractography. Nat Commun, 2017, 8: 1349.

Mak-McCully, R.A., Rolland, M., Sargsyan, A., et al. Coordination of cortical and thalamic activity during non-REM sleep in humans. Nat Commun, 2017, 8: 15499.

Martin, N., Lafortune, M., Godbout, J., et al. Topography of age-related changes in sleep spindles. Neurobiology of Aging, 2013, 34: 468–476.

McCormick, D.A., Bal, T. SLEEP AND AROUSAL: Thalamocortical Mechanisms. Annu. Rev. Neurosci., 1997, 20: 185–215.

Menzler, K., Belke, M., Wehrmann, E., et al. Men and women are different: Diffusion tensor imaging reveals sexual dimorphism in the microstructure of the thalamus, corpus callosum and cingulum. NeuroImage, 2011, 54: 2557–2562.

Piantoni, G., Poil, S.-S., Linkenkaer-Hansen, K., et al. Individual Differences in White Matter Diffusion Affect Sleep Oscillations. Journal of Neuroscience, 2013, 33: 227–233.

Robinson, P.A., Rennie, C.J., Wright, J.J., Bahramali, H., Gordon, E., Rowe, D.L. Prediction of electroencephalographic spectra from neurophysiology. Phys. Rev. E, 2001, 63: 021903.

Romans, S.E., Kreindler, D., Einstein, G., Laredo, S., Petrovic, M.J., Stanley, J. Sleep quality and the menstrual cycle. Sleep Medicine, 2015, 16: 489–495.

Sharkey, K.M., Crawford, S.L., Kim, S., Joffe, H. Objective sleep interruption and reproductive hormone dynamics in the menstrual cycle. Sleep Medicine, 2014, 15: 688–693.

Steriade, M. Corticothalamic resonance, states of vigilance and mentation. Neuroscience, 2000, 101: 243–276.

Takao, H., Hayashi, N., Ohtomo, K. Sex dimorphism in the white matter: Fractional anisotropy and brain size: Sex Dimorphism in the White Matter. J. Magn. Reson. Imaging, 2014, 39: 917–923.

Timofeev, I., Bazhenov, M., Sejnowski, T.J., Steriade, M. Contribution of Intrinsic and Synaptic Factors in the Desynchronization of Thalamic Oscillatory Activity. Thalamus & Related Systems 1, 2001, 53–69.

Tournier, J.-D., Yeh, C.-H., Calamante, F., Cho, K.-H., Connelly, A., Lin, C.-P. Resolving crossing fibres using constrained spherical deconvolution: Validation using diffusion-weighted imaging phantom data. NeuroImage, 2008, 42: 617–625.

Tustison, N.J., Cook, P.A., Klein, A., et al. Large-scale evaluation of ANTs and FreeSurfer cortical thickness measurements. NeuroImage, 2014, 99: 166–179.

Ujma, P.P., Konrad, B.N., Genzel, L., et al. Sleep Spindles and Intelligence: Evidence for a Sexual Dimorphism. J. Neurosci., 2014, 34: 16358–16368.

Veraart, J., Sijbers, J., Sunaert, S., Leemans, A., Jeurissen, B. Weighted linear least squares estimation of diffusion MRI parameters: Strengths, limitations, and pitfalls. NeuroImage, 2013, 81: 335–346.

Wang, S.S.-H., Shultz, J.R., Burish, M.J., et al. Functional Trade-Offs in White Matter Axonal Scaling. Journal of Neuroscience, 2008, 28: 4047–4056.

